# High-throughput nano-flow proteomics using a dual-column electrospray source

**DOI:** 10.1101/2024.02.13.580084

**Authors:** An Staes, Katie Boucher, Sara Dufour, Teresa Mendes Maia, Evy Timmerman, Delphi Van Haver, Jarne Pauwels, Hans Demol, Jonathan Vandenbussche, Francis Impens, Simon Devos

## Abstract

With current trends in proteomics, especially regarding clinical and low input (to single cell) samples, it is increasingly important to both maximize the throughput of the analysis and maintain as much sensitivity as possible. The new generation of mass spectrometers (MS) are taking a huge leap in sensitivity, allowing to analyze samples with shorter liquid chromatography (LC) methods while digging as deep in the proteome. However, the throughput can be doubled by implementing a dual column nano-LC-MS configuration. For this purpose, we used a dual-column setup with a two-outlet electrospray source, and compared it to a classic dual-column setup with a single-outlet source.

## INTRODUCTION

Recent advances in liquid chromatography (LC) and especially mass spectrometry (MS) allow researchers to dig deeper in the proteome of increasingly smaller samples^1,2,3,4^. MS infrastructure is now able to scan analytes eluting from the LC system with incredible speed and sensitivity, using up to nearly a 100% of ions that are entering the MS system^5,6^. This is especially valuable for single cell and (single cell) laser capture microdissected (LCM) tissue proteomics applications, as well as low input clinical proteomics. Such applications, as well as clinical proteomics samples in general often consist of large sample cohorts, benefiting the most of high throughput. An option in increasing analysis throughput is the use of isobaric mass tags^7,8,9,10,11^. Although promising, the limited number of mass tags is still not sufficient for the cohort size of clinical samples or single cell analysis^12,^ and on top suffer from the co-isolation problem^13,14,15^. The new generation of LC and MS systems, in combination with data-independent acquisition (DIA) methods, enables rapid and deep shotgun proteomics in combination with shorter LC runs^16,17^. Even with capillary LC flow instead of nano LC flow^18,19^. Although capillary LC (cLC) is more robust and delivers more throughput, nano LC (nLC) is much more sensitive^18^. Still, nLC suffers from lower throughput and reproducibility, but is the ideal setup for low input proteomics. Another factor that needs to be addressed using nLC, especially with low input samples, is sample carryover^20,21,22^. In order to minimize carryover, cleaning steps are incorporated to eliminate carryover caused by the injection system and the trapping column, and blank samples are run in between analytical runs to further alleviate the issue^20^. These measures are however detrimental for sample throughput since even though using shorter runs, the amount of dead time is higher than the sampling time.

However, the throughput of nLC-MS can be increased without abandoning sample carryover limiting-measures by using a dual pump nLC system. Such system allows for two parallel flow paths, each with its own trapping column and analytical column. While an analytical run is carried out on one flow path, aligned with the inlet of the MS for data acquisition, a blank run is carried out on the second flow path that is directed to the waste as shown before by Kreimer *et al*.^22^, Hosp *et al*.^23^ and Orton *et al*.^24^. By adding a dual source that is able to pneumatically align two separate flow path emitters alternately in-line with the MS inlet, cross contamination between the two flow paths is avoided ^25^. Here we compared two different setups to accomplish a dual column system for high throughput nano-flow proteomics. In the first setup a post-column valve is used that directs the analytical flow path to the MS inlet, and the blank flow path to the waste. The setup is feasible and doesn’t require much extra hardware, but suffers from post-column dead volume leading to loss of chromatographic resolution^24^. In a second setup, a dual-column nLC electrospray source is used (Phoenix S&T), which carries both analytical columns (thermostatted) and uses a pneumatic switching mechanism to align the analytical flow path with the MS inlet, hereby eliminating the abovementioned post-column volume. And in contrast to the dual column setup described by Webber *et al*.^21^, the use of this dual source allows an optimal position of each flow path to the MS inlet, critical for nLC-MS, and assures a safe distance between the two flow paths for eliminating cross-contamination. Moreover, no split flows are used, which are sensitive to clogging and subsequent decreased performance because of the resulting increase in pressure. The advantage compared to Kreimer *et al*.^22^ is that memory on the analytical column is totally avoided.

All modules are commercially available and methods are easily reproduced.

### EXPERIMENTAL SECTION

#### General setup

20 μg of commercial Pierce Hela Protein Digest Standard (Thermo Scientific) was re-dissolved in 400 μl loading solvent A (0.1% trifluoroacetic acid in water/acetonitrile (ACN) (99.5:0.5, v/v)) of which 10 μl was injected for LC-MS/MS analysis on an Ultimate 3000 Pro Flow dual nanoLC system in-line connected to a Q Exactive HF Biopharma mass spectrometer (Thermo Scientific). For each setup 10 replicates were run. In case of tandem setups, 10 replicates of each flow path were run.

#### Analytical LC method

For all setups trapping was performed at 20 μl/min for 2 min in loading solvent A on a 5 mm trapping column (PepMap™ Neo Trap Cartridge; Thermo Scientific). After loading the peptides were further separated on a 250 mm analytical column (made in-house, with either a fritted or in-house pulled needle capillary, 75 μm I.D., 1.9 μm beads, C18 Reprosil-HD; Dr. Maisch) kept at a constant temperature of 45°C in the Ultimate 3000 column oven (only in case of the tandem setup with valve) or in a butterfly oven (Phoenix S&T). Peptides were eluted by a non-linear gradient starting at 0.5% MS solvent B reaching 26% MS solvent B (0.1% FA in acetonitrile) in 30 min, 44% MS solvent B in 38 min, 56% MS solvent B in 40 minutes followed by a 5-minute wash at 56% MS solvent B and re-equilibration with MS solvent A (0.1% FA in water) at a constant flow rate of 250 nl/min.

#### Blank LC method

During the analytical run, at least one blank sample is run on the other column using the second nanoLC pump, next to a cleaning of the autosampler (offline from the rest of the flow path) and a cleaning of the trapping column of the other flow path (offline from the analytical column).

After injecting the blank sample, the analytical column is cleaned with three consecutive gradients, going from 0.5% solvent B from 4 min to 90% solvent B at 10 min, rinsing for 2 min with 90% solvent B, going back to 0.5% Solvent B at 12.1 min and increasing back to 90% solvent B at 18 min for the second gradient. After 2 min of rinsing with 90% solvent B. The third gradient is started at 20.1 min with 0.5% B to be increased to 90% solvent B at 26 min, rinse the last time with 90% solvent B for 2 min and re-equilibrated from 26.1min on with 0.5% solvent B until the end of the run at 60 min. All three gradients are run at 250 nl/min. The trapping column is kept in-line with the analytical column for the first 12 min. After that it switches offline from the analytical column to avoid rinsing the trapping column with a high percentage solvent B and as such avoid the elution of material from the trapping column onto the analytical column.

The trapping itself is then cleaned by two gradients formed by the loading pump at a slightly higher flow rate. Before the trap is switched offline with the analytical column, the autosampler is first cleaned by the loading pump at high flow rate running two gradients (one forward and one reverse). After 4 min the flow rate of the loading pump is increased from 20 μl/min to 100 μl/min and a gradient is started to clean the autosampler. Loading solvent B (0.1% TFA in 80:20 ACN:H2O) is increased from 0% at 4.1 min to 95% loading solvent B at 6 min (forward gradient) to be decreased again to 0% solvent B at 10 min (reverse gradient). At 10.1 min the flow rate of the loading pump is decreased to 30 μl/min. At 12 min, the autosampler is put in-line with the trapping column, and the trapping column is put offline from the analytical column. The loading pump can now perform the cleaning gradient for the trapping column at a flow rate of 30 μl/min. This is started at 10.1 min at 0% loading solvent B and is increased to 95% loading solvent B at 20 min. The trapping is rinsed with 95% loading solvent B until 21 min, after which the flow rate is decreased to 20 μl/min and the loading solvent B to 0%, and kept as such for the remainder of the run.

#### Column switching and ESI parameters

In case of the tandem setup with one outlet, the switching between the two flow paths was controlled by the nano-valve in the autosampler (‘MS switching valve’). A trigger is set at the beginning of each run to either “collect” or “drain” to set the nano-valve into the correct position. In case of the full tandem setup, the source is triggered to both switch the high voltage of the dual source as well as the pneumatic switch of the dual source (Pneu-Nimbus source Phoenix S&T) to put the emitter or column in-line with the inlet of the mass spectrometer. The trigger was performed by enabling a relay trigger in through the autosampler in the method.

Different spray voltage settings were used for each setup and ranged from 2.2 kV to 2.8 kV. In the case of the one outlet setup or fritted column setups, Fossil ionTech emitters were used.

#### MS method

The mass spectrometer was operated in data-independent mode, automatically switching between MS and MS/MS acquisition. Full-scan MS spectra ranging from 375-1500 m/z with a target value of 5E6, a maximum fill time of 50 ms and a resolution at of 60,000 were followed by 30 quadrupole isolations with a precursor isolation width of 10 m/z for HCD fragmentation at an NCE of 30% after filling the trap at a target value of 3E6 for maximum injection time of 45 ms. MS2 spectra were acquired at a resolution of 15,000 at 200 m/z in the Orbitrap analyzer without multiplexing. The non-overlapping isolation intervals with optimal window placement ranging from 400-900 m/z of 10 m/z were created with the Skyline software tool.

In the one column setup the blank samples were recorded as well. Here the mass spectrometer was operated in data-dependent mode, automatically switching between MS and MS/MS acquisition for the 12 most abundant ion peaks per MS spectrum. Full-scan MS spectra (350-1500 m/z) were acquired at a resolution of 120,000 in the Orbitrap analyzer after accumulation to a target value of 3,000,000 with a maximum time of 50 ms. The 12 most intense ions above a threshold value of 90,000 and a charge ranging from 2-5 were isolated with a width of 1.2 m/z for fragmentation at a normalized collision energy of 28% after filling the trap at a target value of 100,000 for maximum 50 ms with a dynamic exclusion of 15s. MS/MS spectra (200-2000 m/z, fixed first mass 145.0 m/z) were acquired at a resolution of 30,000 in the Orbitrap analyzer.

#### Data Analysis

LC-MS/MS runs of all samples were searched together using the DIA-NN algorithm (version 1.8.2 beta 27) in library free mode. Spectra were searched against the human protein database (Swiss-Prot database release version of 2023_08), containing 20423 sequences (www.uniprot.org). Enzyme specificity was set as C-terminal to arginine and lysine, also allowing cleavage at proline bonds with a maximum of two missed cleavages. Variable modifications were set to oxidation of methionine residues and acetylation of protein N-termini with a maximum of 5 variable modifications and only cysteine carbamidomethylation was set as fixed modification. Mainly default settings were used, except for the addition of a 400-900 m/z precursor mass range filter and minimum precursor charge of 2.

Blank runs were searched separately with Comet (v2023.01 rev.2)^26,27^ starting directly from the raw file. against the human protein database (Swiss-Prot database release version of 2023_08), containing 20423 sequences (www.uniprot.org). Enzyme specificity was set as C-terminal to arginine and lysine, also allowing cleavage at proline bonds with a maximum of two missed cleavages. Variable modifications were set to oxidation of methionine residues and acetylation of protein N-termini and only cysteine carbamidomethylation was set as fixed modification. Tolerance was set to 0.02 m/z. Results were filtered on a e-value of 0.01.

## RESULTS AND DISCUSSION

Two dual-column setups were configured on the same dual-pump LC system, coupled to the same MS instrument. In a first setup the two flow paths (with separate trapping and analytical column) are connected to a single electrospray needle through a connection capillary (Figure 1A). A 6-port nanovolume valve (100 μm port) between the fritted columns and the connection capillary is directing the analytical flow path to the MS inlet, and the blank flow path to the waste. In the second setup, the post-column switching valve is omitted through the use of a dual-column nLC electrospray source (Pneu-Nimbus source, Phoenix S&T) with an integrated, programmable pneumatic switch to align the analytical (emitter) column to the MS inlet (Figure 1B). Because the columns are placed outside of the LC system’s column compartment, a ‘butterfly’ column heater is used on-source (Butterfly heater, Phoenix S&T).

**Figure 1.**
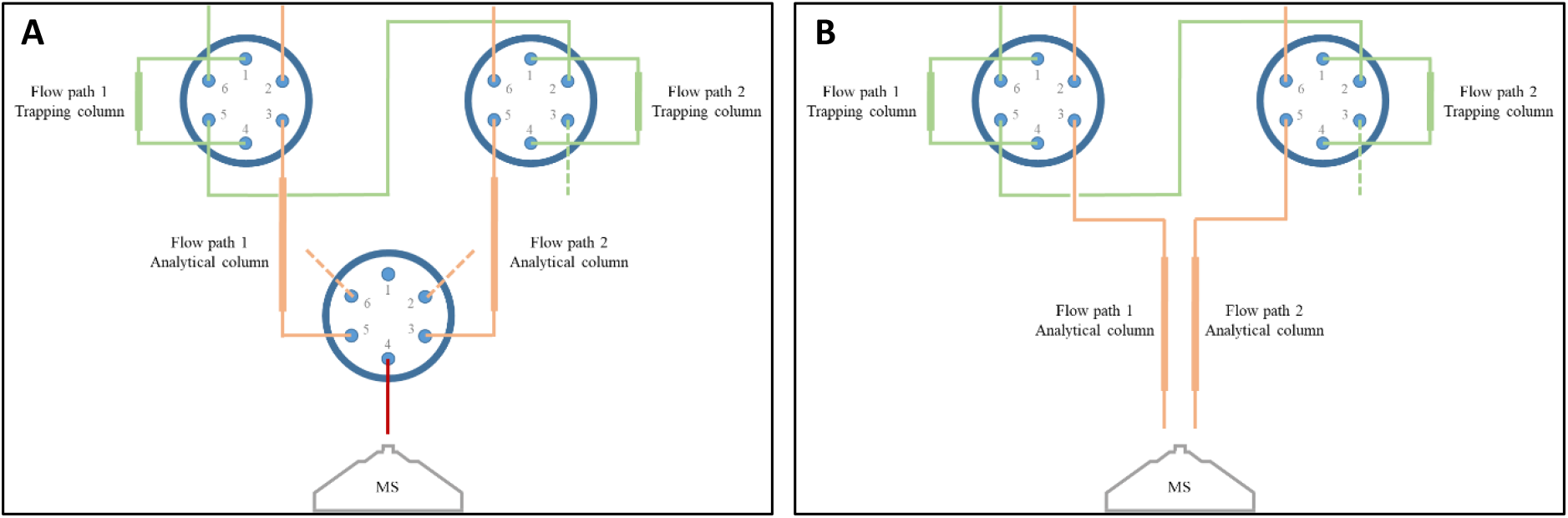
Configuration of the two dual-column setups. A. Single-outlet dual-column setup. A sample can be trapped on flow path 1 (trap valve position flow path 1 to 6-1), while the analytical column of flow path 1 is directed to the MS inlet (MS valve position to 6-1). After switching the trap valve of flow path 1 (position 1-2) the sample is separated and measured with MS. Meanwhile, a blank sample can be loaded on the trapping column of flow path 2 (trap valve position flow path 1 to 1-2, trap valve position flow path 2 to 1-2). After switching the trap valve of flow path 2 (position 6-1), a blank run is performed on the analytical column of flow path 2, directed to the waste ((MS valve position to 6-1). In the next run, a sample is loaded on the trapping column of flow path 2 and measured, while a blank is loaded on the trapping column of flow path 1. B. Two-outlet dual-column setup. The same configuration is used as with the single-outlet dual-column setup, but the two columns are placed in a column heated located on a dual-column electrospray source, hereby omitting the need for a MS valve.

Using both dual-column setups, ten HeLa protein digest standards (Pierce) were alternately separated on each flow path (A and B), using a one-hour LC gradient and eluting peptides analyzed with DIA-MS on a Orbitrap Q Exactive HF. Blanks were run alternately on the off-line flow path, using a multi-gradient cleaning method for rinsing the autosampler, the trapping column and the analytical column.

With the single-outlet dual-column setup (Figure 1A), an average of 2646 and 2392 proteins were identified, on flow path A and B, respectively (Figure 2A). However, with the two-outlet dual-column setup (Figure 1B), the average number of proteins identified increased to 5167 and 5196, on flow path A and B, respectively (Figure 2A). The number of identifications is roughly doubled with the two-outlet setup, and is in large due to post-column volume and accompanied LC peak broadening of the single outlet setup. The average FWHM with the two-outlet setup is 0.12 min and 0.13 min, on flow path A and B, respectively, as compared to 0.24 min and 0.28 min with the single-outlet setup (Figure 2B).

**Figure 2.**
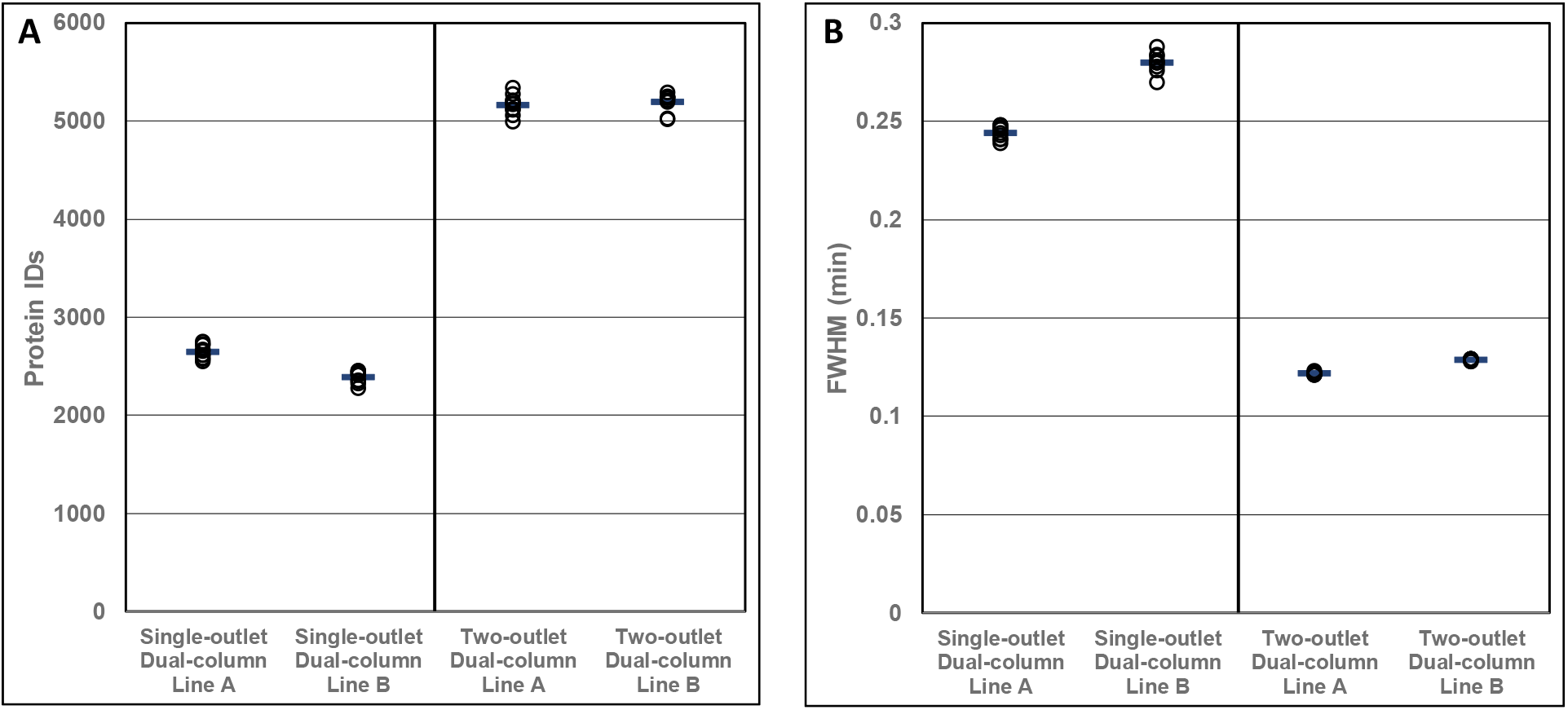
Comparison of the number of protein identifications (left panel) and peak width (FWHM; right panel) using both dual column setups, on both flow paths.

Different in-house produced columns were used for testing both setups, i.e. fritted columns for the single-outlet setup and emitter columns for the two-outlet setup. In order to exclude any differences in performance between the two column types, the fritted columns used in the dual flow path configuration were also tested in a single flow path configuration (placed in the butterfly heater close to the source). The number of identifications resulting from these single column setups was in the same range as in the dual column setup. Although the dual column setup with emitter columns (Figure 1B) remains the best performing configuration, using a fritted column either in a single column setup (placed close to the source in the butterfly heater) or in dual column setup (on dual source using electrospray needles) shows better results than the dual column setup using the MS switching valve (Figure 1A) (Supplementary Figure 1). This shows that the peak broadening, and hence loss of protein IDs, is largely due to the MS switching valve. We also see a lower reproducibility when using fritted columns with electrospray needles as compared to emitter columns (Supplementary Figure 1). This will be due to the fact that an extra connection is needed, most probably leading to the higher variability by possible introducing dead volume.

Next to the increased number of identifications, the CV dropped from 5% to 1.9% by using the dual outlet setup, showcasing the reproducibility of the different flow paths and columns (Supplementary Table 1). The CV of the in-needle tandem setup is in the same range as a single column setup (1.86% for the tandem in needle setup versus 1.09% for the single column setup) pointing out the reproducibility of the two flow paths. Although applying a nanoflow setup, this CV is much lower compared to the 18% of the dual column setup of Webber *et al*.^21^ and 20% of the dual trap single column setup (micro flow) of Kreimer *et al*.^22^. By running the tandem setup, the duty cycle (i.e. the percentage of the total run time where actual sample data is generated) of each run is increased from 24% for a single column setup to 51% for the dual column setup (Supplementary Table 2).

Analysis of the blank runs shows the necessity of implementing cleaning runs. But the main memory factor seems to be the trapping column, derived from the fact that the only remaining carry-over identifications were seen in the first cleaning gradient where the trapping column is in-line with the analytical column (Supplementary Table 3). This is in accordance to Kreimer *et al*.^22^ and Bach *et al*.^19^, who are using different trapping columns in combination with the same analytical column, and without blank runs in between. Compared to the dual-trap, single column setup, the dual column setup has the advantage of enabling a cleaning of the autosampler loop with organic solvent but without eluting leftover sample onto the trapping column, assuring no cross-contamination between samples. A cleaning of the analytical column is done in several gradients as well to be more confident about minimizing memory. Indeed, in single cell analysis for example, the smallest trace of memory can be detrimental.

In conclusion, in order to increase throughput in (low input) nano-LC-MS/MS and minimizing sample carry over, while safeguarding the proteome coverage depth and the run-to-run reproducibility, using a dual emitter column setup with a dual-column nLC electrospray source is the best solution.

## ASSOCIATED CONTENT

### Supporting Information

Supplementary Figure 1. Number of protein IDs from all setups

Supplementary Table 1. CV values from different setups

Supplementary Table 2. Duty cycle values

Supplementary Table 3. Number of protein IDs in blank runs

## AUTHOR INFORMATION

## ACKNOWLEDGEMENTS

The authors would like to thank Sau Lan Staats from Phoenix S&T and Joerg Niebel from MS Wil for their technical assistance in installing and configuring the Nimbus Dual Column Source.

## Notes

### Competing Interest Statement

The authors have declared no competing interest.

