## Supplementary Information for "High-throughput nano-flow proteomics using a dual-column electrospray source"

### SUPPLEMENTARY FIGURES

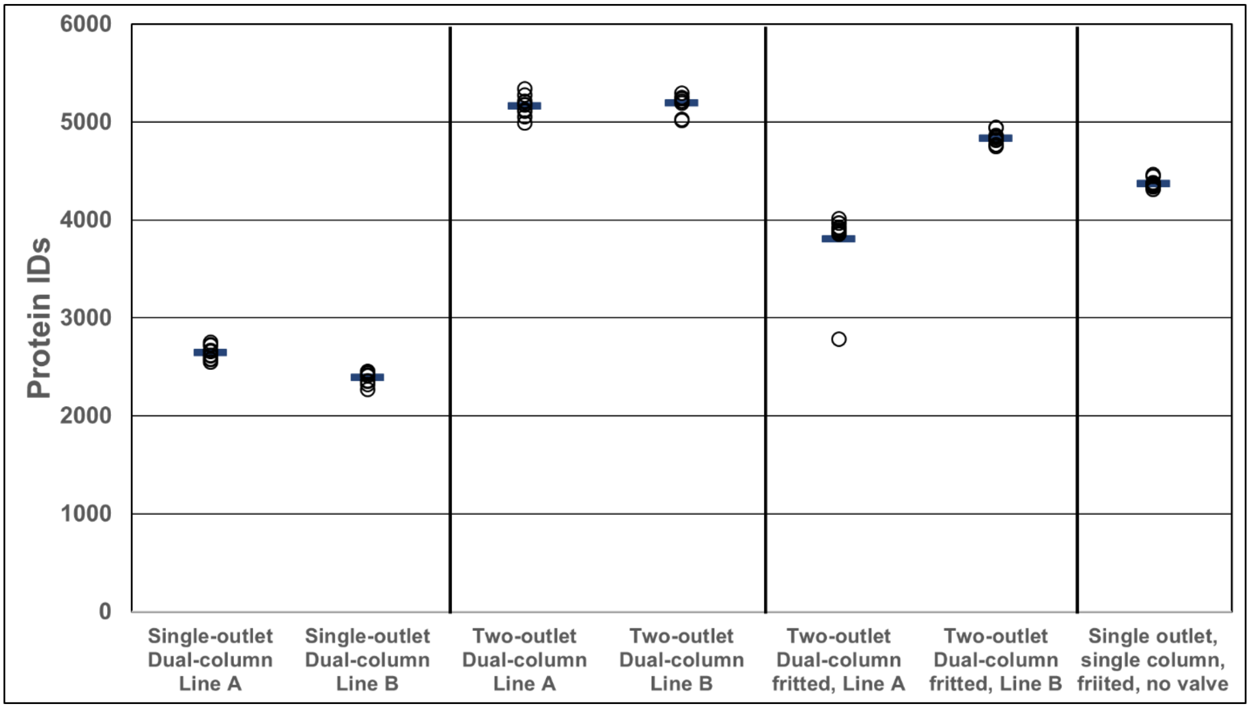


**Supplementary Figure 1.** Comparison of the number of protein identifications using both single and dual column setups, and separate both flow paths.

**Supplementary Table 1.** Calculated CV values of both single and dual column setups, and separate both flow paths.

| | Average | Median | StDev | CV |
|---|---|---|---|---|
| Single-outlet Dual-column Line A | 2645.8 | 2637.5 | 70.4884545 | 2.66% |
| Single-outlet Dual-column Line B | 2391.5 | 2413.5 | 61.8820563 | 2.59% |
| Average | 2518.65 | 2502.5 | 145.552405 | **5.78%** |
| Two-outlet Dual-column Line A | 5166.9 | 5175.5 | 100.983442 | 1.95% |
| Two-outlet Dual-column Line B | 5195.8 | 5238.5 | 94.9149327 | 1.83% |
| Average | 5181.35 | 5210 | 96.5277571 | **1.86%** |
| Two-outlet Dual-column, fritted, Line A* | 3809.8 | 3920 | 51.771131 | 1.36% |
| Two-outlet Dual-column, fritted, Line B | 4835.9 | 4830 | 67.7896256 | 1.40% |
| Average | 4403.78947 | 4749 | 471.678867 | **10.71%** |
| Single outlet Single column, fritted, no valve | 4374.1 | 4360.5 | 47.806206 | **1.09%** |

**Two-outlet Dual-column fritted, Line A: outlier value of 2785 was left out*

**Supplementary Table 2.** Calculated duty cycle values of both single and dual column setups, and separate both flow paths.

| | | min RT (min) | max RT (min) | Duty Cycle (%) | Duty Cycle, Blank incl (%) |
|---|---|---|---|---|---|
| **Single Column** | Hela run 1 | 16.64 | 45.76 | 48.54 | 24.27 |
| | Hela run 2 | 16.70 | 45.75 | 48.41 | 24.21 |
| | Hela run 3 | 16.63 | 45.70 | 48.44 | 24.22 |
| | Hela run 4 | 16.70 | 45.70 | 48.33 | 24.17 |
| | Hela run 5 | 16.63 | 45.66 | 48.38 | 24.19 |
| | Hela run 6 | 16.59 | 45.63 | 48.40 | 24.20 |
| | Hela run 7 | 16.56 | 45.62 | 48.43 | 24.22 |
| | Hela run 8 | 16.56 | 45.63 | 48.45 | 24.22 |
| | Hela run 9 | 16.49 | 45.63 | 48.56 | 24.28 |
| | | | | **Average** | **24.22** |
| **Dual Column Line A** | Hela run 1 | 13.21 | 45.63 | 54.03 | 54.03 |
| | Hela run 2 | 13.40 | 45.62 | 53.70 | 53.70 |
| | Hela run 3 | 13.84 | 45.52 | 52.80 | 52.80 |
| | Hela run 4 | 13.63 | 45.46 | 53.05 | 53.05 |
| | Hela run 5 | 13.56 | 45.41 | 53.09 | 53.09 |
| | Hela run 6 | 13.34 | 45.39 | 53.43 | 53.43 |
| | Hela run 7 | 13.68 | 45.36 | 52.79 | 52.79 |
| | Hela run 8 | 13.27 | 44.71 | 52.39 | 52.39 |
| | Hela run 9 | 13.21 | 45.34 | 53.56 | 53.56 |
| | | | | **Average - Line A** | **53.20** |
| **Dual Column Line B** | Hela run 1 | 16.51 | 46.57 | 50.09 | 50.09 |
| | Hela run 2 | 16.51 | 46.56 | 50.07 | 50.07 |
| | Hela run 3 | 16.48 | 46.58 | 50.17 | 50.17 |
| | Hela run 4 | 16.48 | 46.04 | 49.27 | 49.27 |
| | Hela run 5 | 16.51 | 45.62 | 48.53 | 48.53 |
| | Hela run 6 | 16.51 | 45.94 | 49.05 | 49.05 |
| | Hela run 7 | 16.41 | 45.68 | 48.78 | 48.78 |
| | Hela run 8 | 16.83 | 45.52 | 47.83 | 47.83 |
| | Hela run 9 | 16.42 | 45.95 | 49.22 | 49.22 |
| | | | | **Average - Line B** | **49.22** |
| | | | | **Average - Line A + B** | **51.21** |

**Supplementary Table 3.** Number of PSMs, peptides and protein identifications in the blank runs (carry-over) of the single column setup.

| | | **Blank before first sample** | **Blank 1** | **Blank 2** | **Blank 3** | **Blank 4** | **Blank 5** | **Blank 6** | **Blank 7** | **Blank 8** | **Blank 9** | **Blank 10** |
|---|---|---|---|---|---|---|---|---|---|---|---|---|
| **Total** | PSMs | 0 | 86 | 84 | 80 | 85 | 74 | 89 | 85 | 86 | 89 | 96 |
| | Peptides | 0 | 82 | 80 | 79 | 82 | 72 | 84 | 81 | 82 | 86 | 93 |
| | Proteins | 0 | 57 | 58 | 53 | 59 | 50 | 58 | 57 | 54 | 55 | 65 |
| **1st Gradient** | PSMs | 0 | 86 | 84 | 80 | 85 | 74 | 89 | 85 | 86 | 89 | 96 |
| | Peptides | 0 | 82 | 80 | 79 | 82 | 72 | 84 | 81 | 82 | 86 | 93 |
| | Proteins | 0 | 57 | 58 | 53 | 59 | 50 | 58 | 57 | 54 | 55 | 65 |
| **2nd Gradient** | PSMs | 0 | 0 | 0 | 0 | 0 | 0 | 0 | 0 | 0 | 0 | 0 |
| | Peptides | 0 | 0 | 0 | 0 | 0 | 0 | 0 | 0 | 0 | 0 | 0 |
| | Proteins | 0 | 0 | 0 | 0 | 0 | 0 | 0 | 0 | 0 | 0 | 0 |
| **3rd Gradient** | PSMs | 0 | 0 | 0 | 0 | 0 | 0 | 0 | 0 | 0 | 0 | 0 |
| | Peptides | 0 | 0 | 0 | 0 | 0 | 0 | 0 | 0 | 0 | 0 | 0 |
| | Proteins | 0 | 0 | 0 | 0 | 0 | 0 | 0 | 0 | 0 | 0 | 0 |
